# Rigbox: an Open-Source Toolbox for Probing Neurons and Behavior

**DOI:** 10.1101/672204

**Authors:** Jai Bhagat, Miles J. Wells, Andrew Peters, Kenneth D Harris, Matteo Carandini, Christopher P Burgess

## Abstract

Setting up an experiment in behavioral neuroscience is a complex process that is often managed with ad hoc solutions. To streamline this process we developed Rigbox, a high-performance, open-source software toolbox that facilitates a modular approach to designing experiments (github.com/cortex-lab/Rigbox). Rigbox simplifies hardware I/O, synchronizes data streams from multiple sources, communicates with remote databases, and implements visual and auditory stimuli presentation. Its main submodule, Signals, allows intuitive programming of behavioral tasks. Here we illustrate its function with two interactive examples: a human psychophysics experiment, and the game of Pong. We give an overview of the other packages in Rigbox, provide benchmarks, and conclude with a discussion on the extensibility of the software and comparisons with similar toolboxes. Rigbox runs in MATLAB, with Java components to handle network communication, and a C library to boost performance.

## Introduction

In behavioral neuroscience, much time is spent setting up hardware and software and ensuring compatibility between them. Experiments often require configuring disparate software to interface with distinct hardware, and integrating these components is no trivial task. Furthermore, there are often separate software components for designing a behavioral task, running the task, and acquiring, processing, and logging the data. This requires learning the fundamentals of different software packages and how to make them communicate appropriately.

Consider a typical experiment focused on decision-making, in which a subject chooses a stimulus amongst a set of possibilities and obtains a reward if the choice was correct (Carandini and Churchland, 2013). The software set-up for this experiment may seem simple: ostensibly, all that is required is software to run the behavioral task, and software to handle the experimental data. However, when considering implementation details for these two types of software, the set-up can grow quite complex. For example, in a typical variant of the Burgess Steering Wheel Task, a mouse must move a steering wheel left or right to choose between two different visual stimuli (Figure 1) (Burgess et al., 2017; Steinmetz et al., 2018). For each trial, if the mouse chooses the correct stimulus, it receives a water reward from a spout. During the experiment, the mouse’s actions are recorded with a body camera and lick detector, and brain activity is recorded with electrodes and manipulated with a laser. Running the task requires software for starting, stopping, and transitioning between task states, presenting the stimuli, and triggering the reward spout and laser. Handling experimental data requires software for acquiring, processing, and logging stimulus history, response history (from the wheel and a lick detector), and subject physiology (from the body camera and the electrodes), and transferring data between servers and databases.

**Figure 1:**
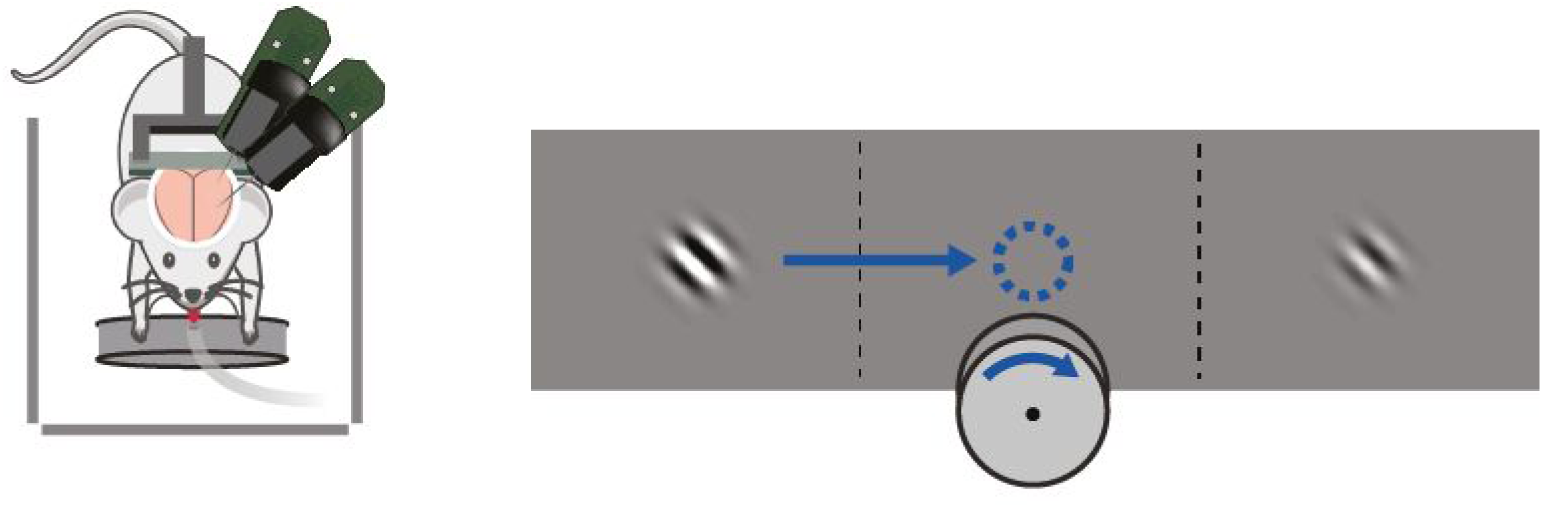
In this experiment, in addition to the software that runs the behavioral task, software is also required to 1) record time-series data from the electrodes, steering wheel, body camera, and lick detectors; 2) trigger outputs to the spout and laser; and 3) transfer data between the rig computer, a remote server (for loading experiment parameters), and a remote database (for saving experiment metadata). (Adapted with permission from Steinmetz et. al 2018)

To address this variety of needs in a single software toolbox, we designed Rigbox (github.com/cortex-lab/Rigbox). Rigbox is modular, high-performance, open-source software for implementing behavioral neuroscience experiments and acquiring experiment-related data. Rigbox facilitates recording, synchronizing, and managing data from a variety of sources. Furthermore, Rigbox promotes bespoke behavioral task design via a framework called Signals, which exploits both object-oriented and functional reactive programming paradigms to allow an experimenter to intuitively define and parameterize an experiment.

## Methods and Results

We begin by giving a general overview of Rigbox. We go on to describe Signals, the core package of Rigbox, and provide two interactive examples of its use: a simple experiment in visual psychophysics, and the game of Pong. We then briefly describe the other packages in Rigbox, and provide benchmarking results.

### Overview

Rigbox is made up of a number of packages which run on two computers, referred to as the “Master Computer” (MC) and “Stimulus/Slave Computer” (SC) (Figure 2). MC is responsible for selecting, parameterizing, starting, and monitoring an experiment via a MATLAB GUI. SC is responsible for running an experiment on a rig and interacting with that rig’s hardware during runtime. MC can control multiple SCs simultaneously.

**Figure 2:**
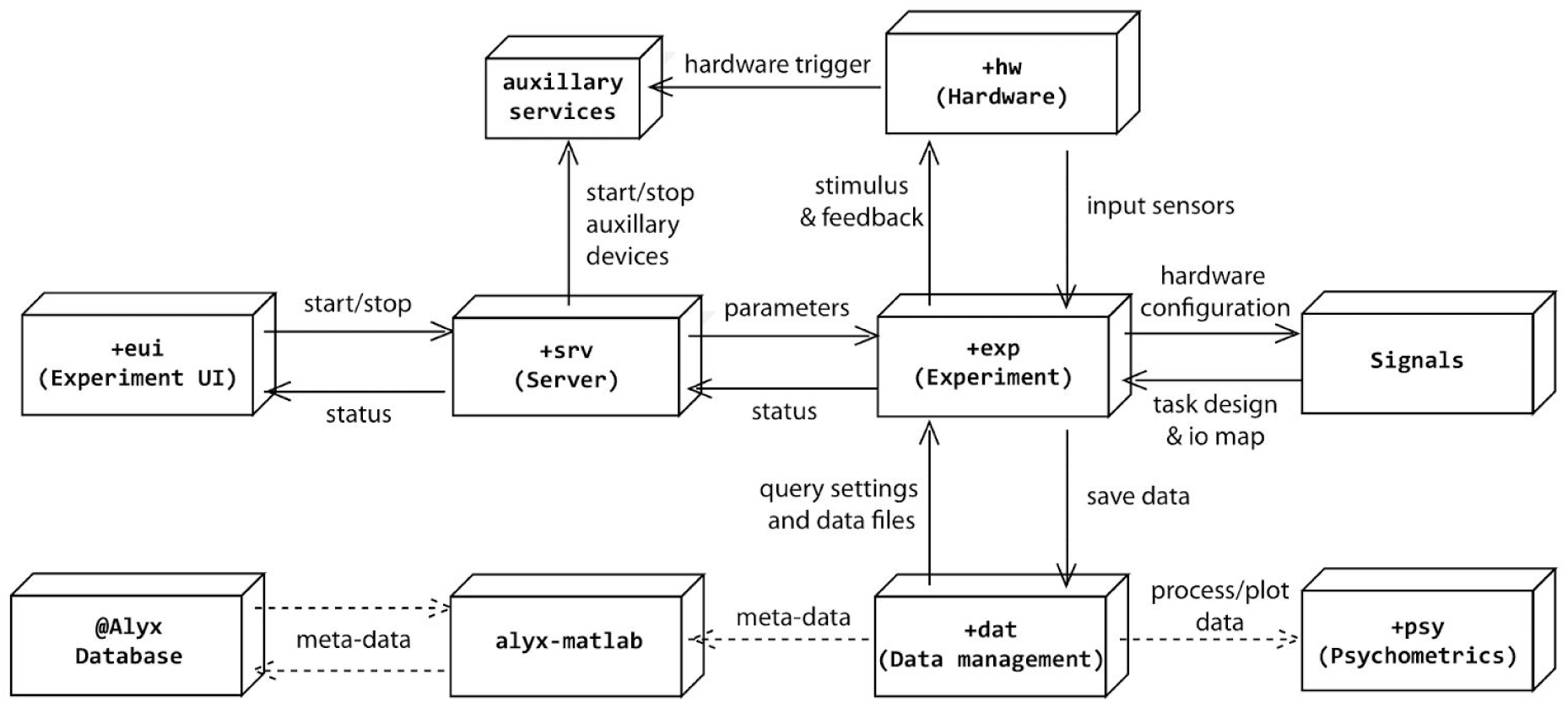
Schematic of Rigbox package interactions. The solid lines represent necessary communication between Rigbox packages, and the dashed lines represent optional communication (for saving data to a remote Alyx database, and for processing/plotting data). The *+eui* package runs a GUI on the master computer (MC), and the *+srv* package launches stimulus presentation on the stimulus computer (SC). Though this figure shows only one direct MC-SC connection, MC can control multiple SCs simultaneously.

MC and SC communicate during runtime via Java WebSockets using TCP/IP. Therefore, it is necessary for both computers to be connected to high-speed internet. The precise computer hardware requirements for SC depend on the complexity of the experiment, and for MC depend on the number of experiments run concurrently (i.e. number of active SCs controlled). For most experiments, typical modern desktop computers running Windows will suffice. SC also requires input/output device(s) for polling hardware inputs and triggering hardware outputs, and optionally requires graphics and sound cards, depending on the complexity of the stimuli to be presented.

Instructions for installation and configuration can be found in the README file and the docs/setup folder of the GitHub repository. This includes information on required dependencies, setting data repository locations, configuring hardware, and setting up communication between the MC and SC computers.

### Signals

Signals was designed for building bespoke behavioral tasks. The framework is built around the paradigm of functional reactive programming, which simplifies problems that deal with change over time (Lew, 2017). Signals represents an experiment as a reactive network whose nodes (“signals”) represent experimental parameters. These signals can evolve over time through interactions with each other. The framework provides a set of input signals which represent time, experiment epochs, and hardware input devices, and a set of output signals which represent hardware output devices. Thus, an entire experiment can simply be thought of as a network which maps hardware inputs to hardware outputs via a set of experimenter-defined transformations (Figure 3).

**Figure 3:**
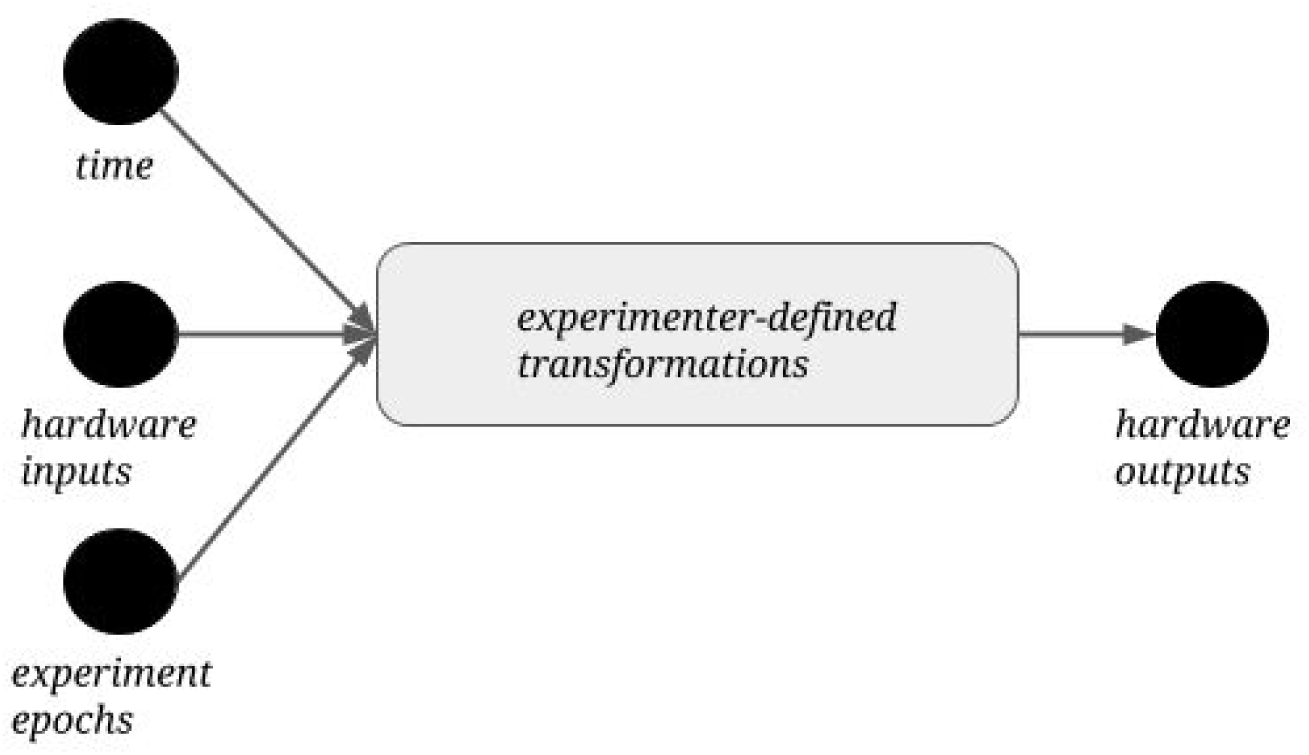
A Signals representation of an experiment. There are three types of input signals in the network, representing time, hardware inputs (such as an optical mouse, keyboard, rotary encoder, lever, etc.), and experiment epochs (such as trial and experiment start and end conditions). The experimenter defines transformations that create new signals (not shown) from these input signals, which ultimately drive hardware outputs (such as a reward valve, blow gun, galvanometer, etc.).

The core goal of Signals is to represent the relationship between experimental parameters with straightforward, self-documenting operations. For example, to define the temporal frequency of a visual stimulus - for example, a drifting grating - an experimenter could create a signal which changes the grating’s phase as a function of time (Figure 4). This is shown in the code below:

**Figure 4:**
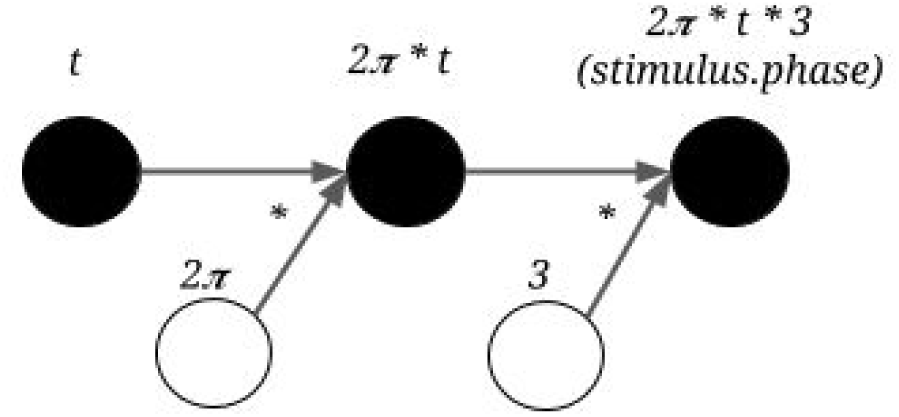
Representation of the time-dependent phase of a visual stimulus in Signals. An unfilled circle represents a constant value - it becomes a node in the network when combined with another signal in an operation (in this instance, via multiplication).

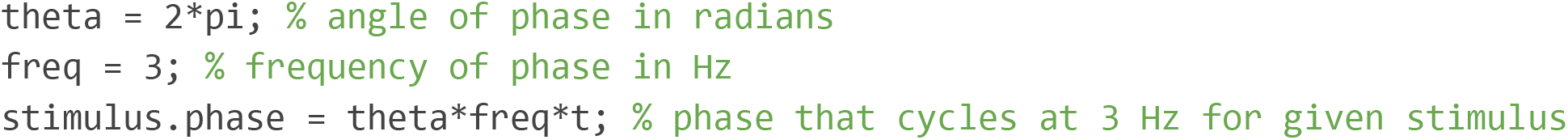

The operations that can be performed on signals are not just limited to basic arithmetic. A number of built-in MATLAB functions (including logical, trigonometric, casting, and array operations) have been overloaded to work on signals as they would on basic numeric or char types. Furthermore, a number of classical functional programming functions (e.g. “map”, “scan”, etc.) can be used on signals. These endow signals with memory, and allow them to gate, trigger, filter, and accumulate other signals (Figure 5).

**Figure 5:**
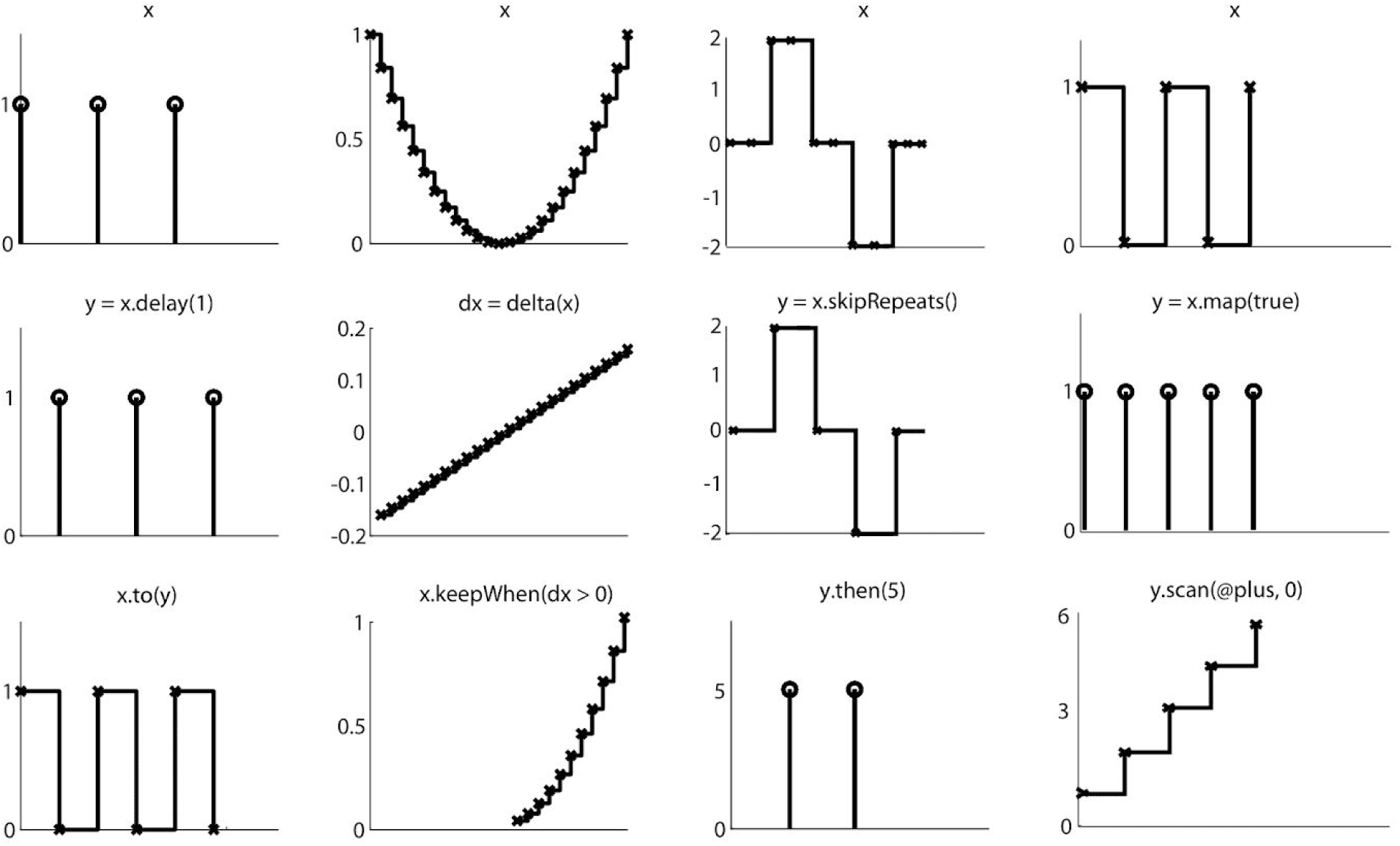
The creation of new signals via example signals methods. Conceptually, a signal can be thought of as both a continuous stream of discrete values, and as a discrete representation whose value changes over time. Each panel represents a signal. The x-axis represents time, and the y-axis represents the signal’s value. Each column depicts a set of related transformations. The second row depicts a signal which results from applying an operation on the signal in the same column’s first row. The third row depicts a signal which results from applying an operation on the signals in the same column’s first and second rows.

With this powerful framework, an experimenter can easily define complex relationships between input and output devices (or more abstractly, between stimuli, response and reward) in order to create a complete experiment protocol. This protocol takes the form of a user-written MATLAB function, which we refer to as an “experiment definition” (“exp def”). Signals runs an experiment by loading this exp def into a network and posting values to the network’s input signals on every iteration of a while loop, which triggers asynchronous propagation through the reactive network. The experiment ends when the “experiment stop” signal is updated (e.g. when a number of correct trials is reached or when the experimenter clicks the “end” button in the MC GUI).

The following is a brief overview of the structure of an exp def. An exp def takes up to seven input arguments:

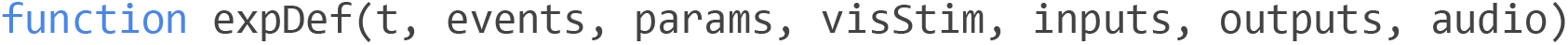

In order, these are 1) the time signal; 2) an events structure containing signals which define the experiment’s epochs, and a set of user-chosen signals to be logged from those defined within the exp def; 3) a parameters structure to define session- or trial-specific signals whose values can be changed directly from the MC GUI before starting an experiment -- parameter defaults are set within the exp def and parameter sets can be saved and loaded across subjects and experiments; 4) the visual stimuli handler which contains as fields all signals which parametrize the display of visual stimuli -- any visual stimulus signal can be assigned various elements (which the viewing model allows to be defined in visual degrees) for being rendered to a screen, and a visual stimulus can be loaded directly from a saved image file; 5) an inputs structure containing signals which map to hardware inputs devices; 6) an outputs structure containing signals which map to hardware output devices; 7) the audio stimuli handler which can contain as fields signals which map to available audio devices.

Tutorials on creating an exp def, examples of working exp defs and standalone scripts (including those mentioned in this paper), and an in-depth overview of Signals can be found in the `signals/docs` folder within the Rigbox repository. Though running a Signals experiment in Rigbox typically requires two computers, the following examples can be run from a single Windows PC, as their only required hardware devices are an optical mouse and keyboard. Readers are encouraged to run these examples upon installing Rigbox and its necessary dependencies.

### Example 1: A Psychophysics Experiment

Our first example of a human-interactive Signals experiment is a script that recreates a psychophysics experiment to study the mechanisms that underlie the discrimination of a visual stimulus (Ringach 1998). In this experiment, the observer looks at visual gratings (Figure 6a) that change rapidly and randomly in orientation and phase. The gratings change so rapidly that they summate in the visual system, and the observer tends to perceive two or three of them as superimposed. The task of the observer is to hit the “ctrl” key whenever the grating’s orientation is vertical. At key press, the probability of detection is plotted as a function of stimulus orientation in the recent past. Typically, this exposes a center-surround type of organization, with orientations near vertical eliciting responses, but orientations further away suppressing responses (Figure 6b). The Signals network representation of this experiment is shown in Figure 7.

**Figure 6:**
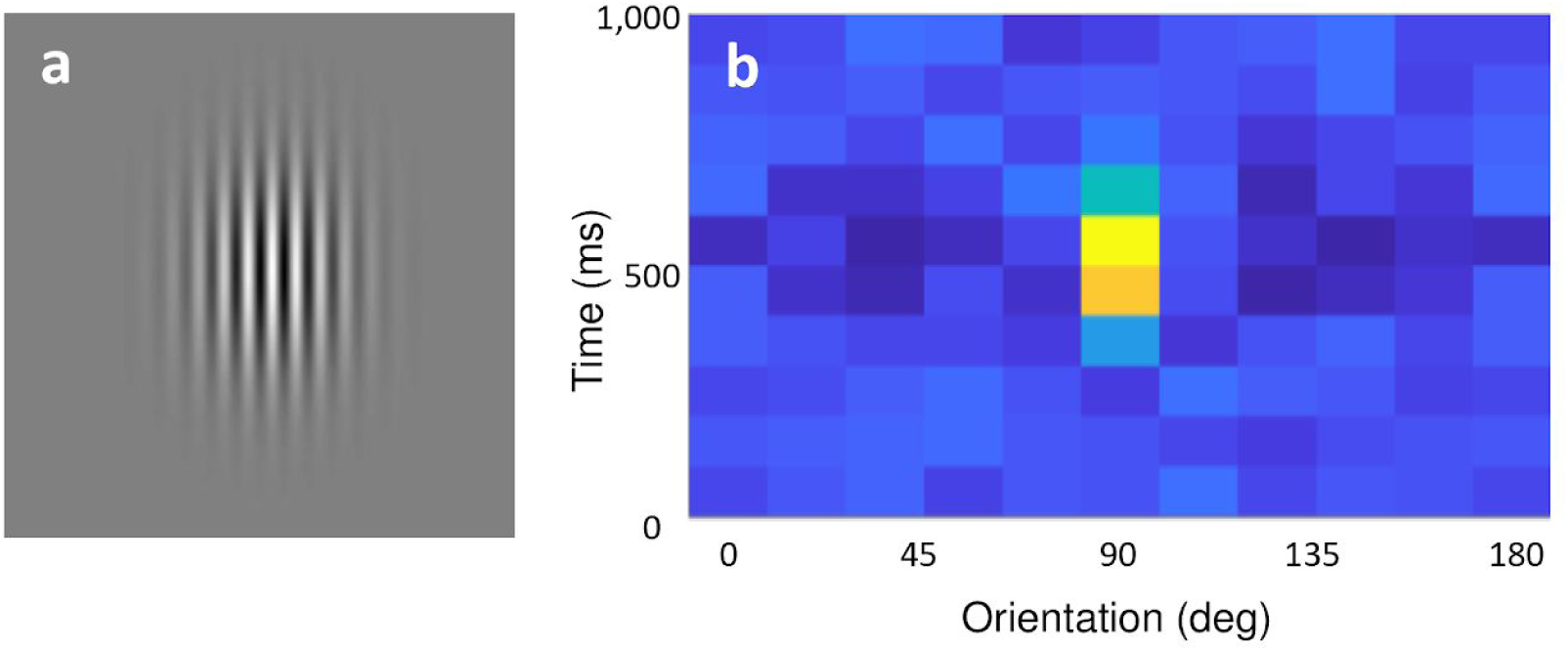
**a)** A sample grating for which the subject is required to respond to via a “ctrl” key press. **b)** A heatmap showing the grating orientations for the ten frames immediately preceding a “ctrl” key press, summed over all “ctrl” key presses for the duration of the experiment. After a few minutes, the distribution of orientations over time at a “ctrl” key press resembles a 2D Mexican Hat wavelet, centered on the orientation the subject was reporting at the subject’s average reaction time. In this example, the subject was reporting a vertical grating orientation (90 degrees) with an average reaction time of roughly 600ms.

**Figure 7:**
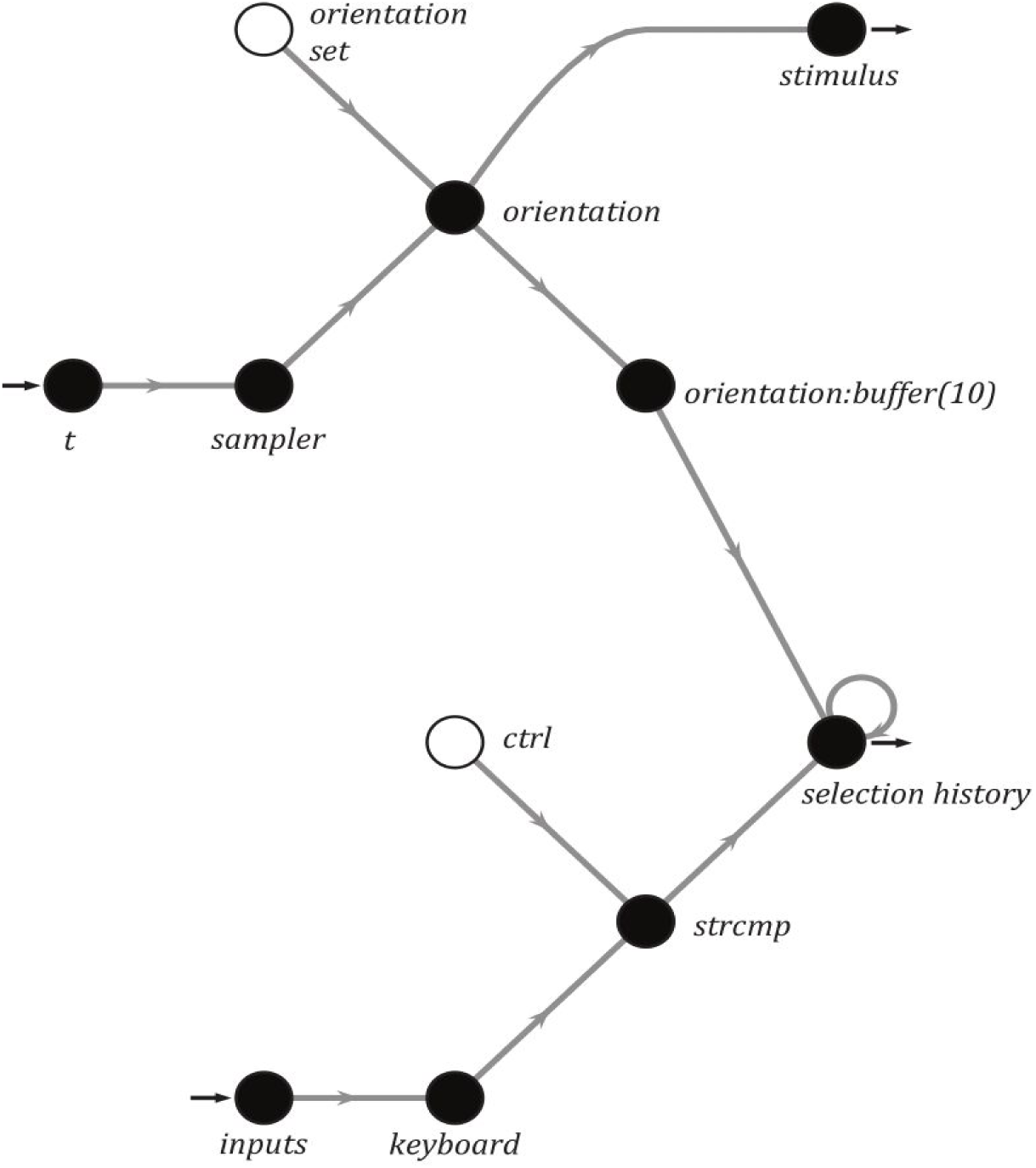
A simplified Signals network diagram of the Ringach experiment.

To run this experiment, simply run the file `signals/docs/examples/ringach98.m` in the Rigbox repository. Below is a breakdown of the thirty lines of code:

First, some constants are defined:

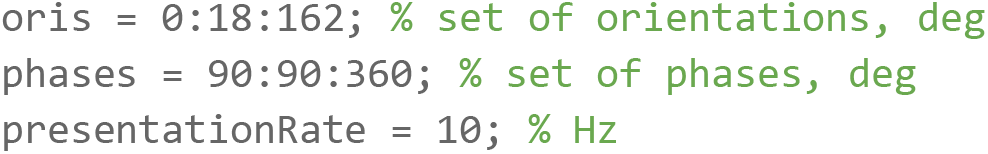
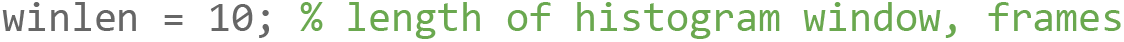

Next, we create a figure and our Signals network:

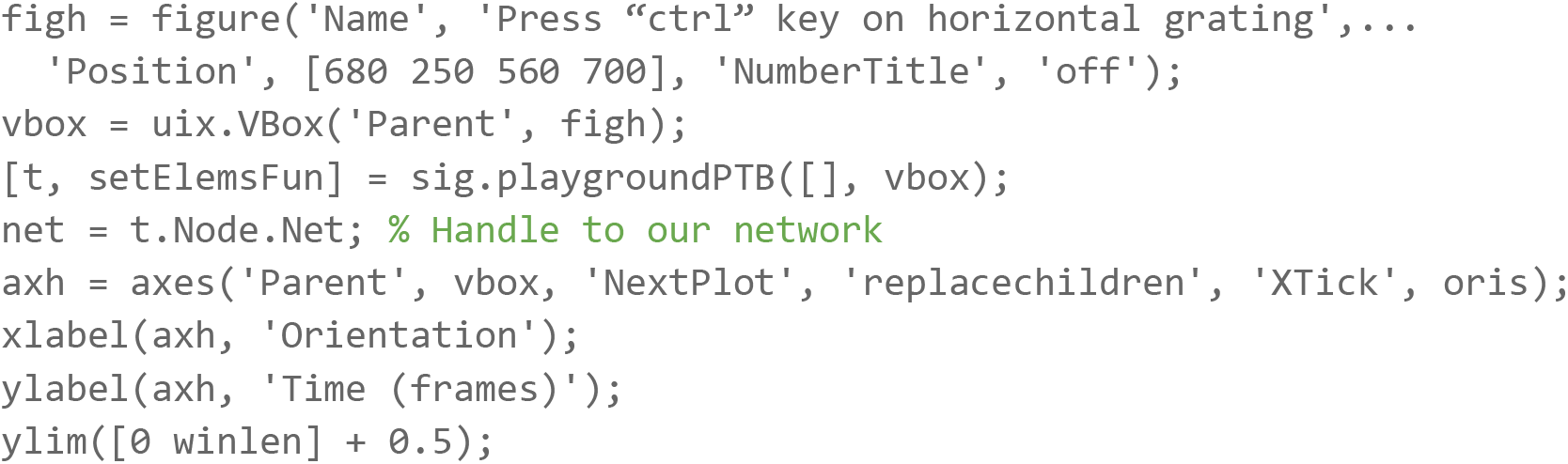

Then, we wire our network:

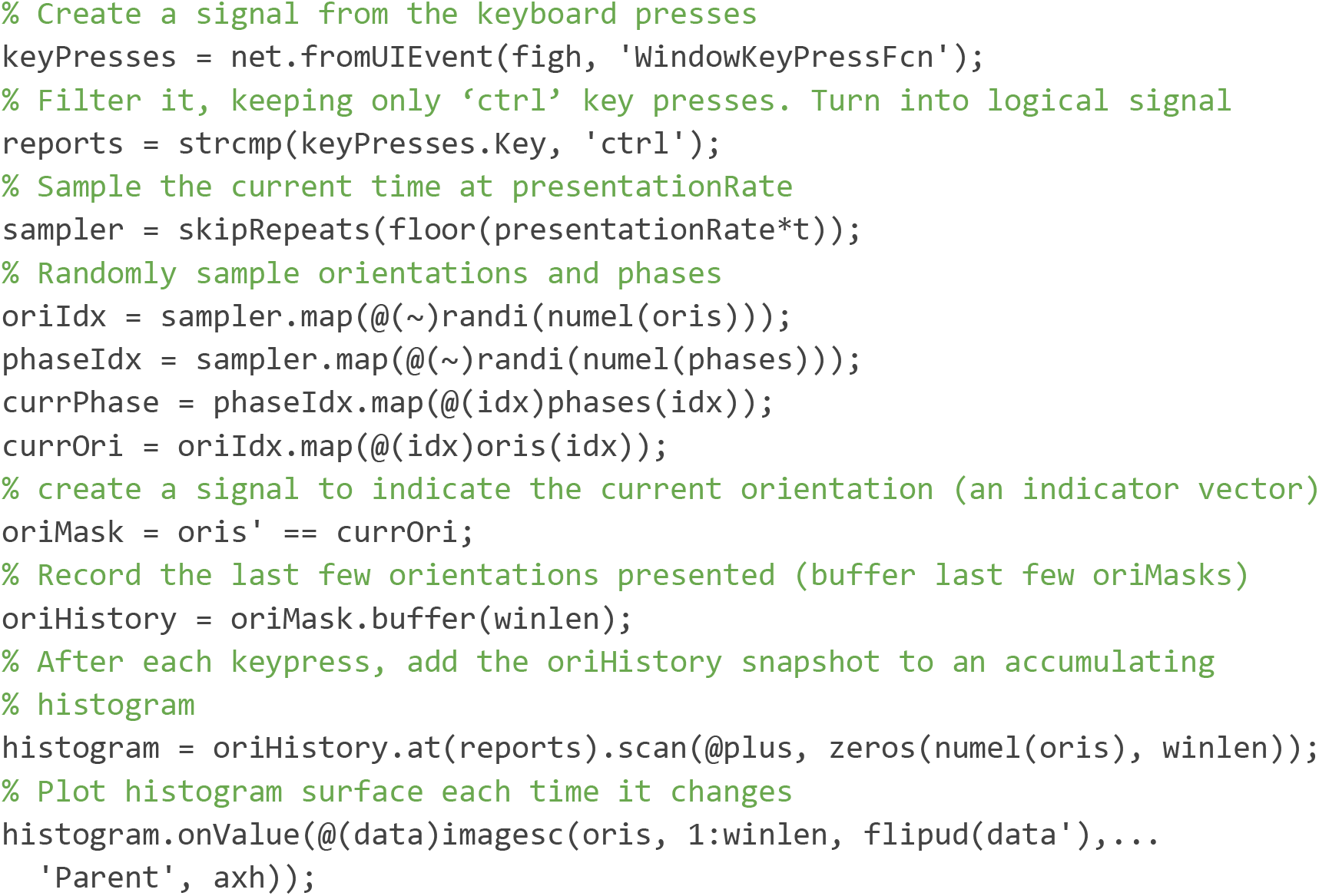

Finally, we create the visual stimulus and send it to the renderer:

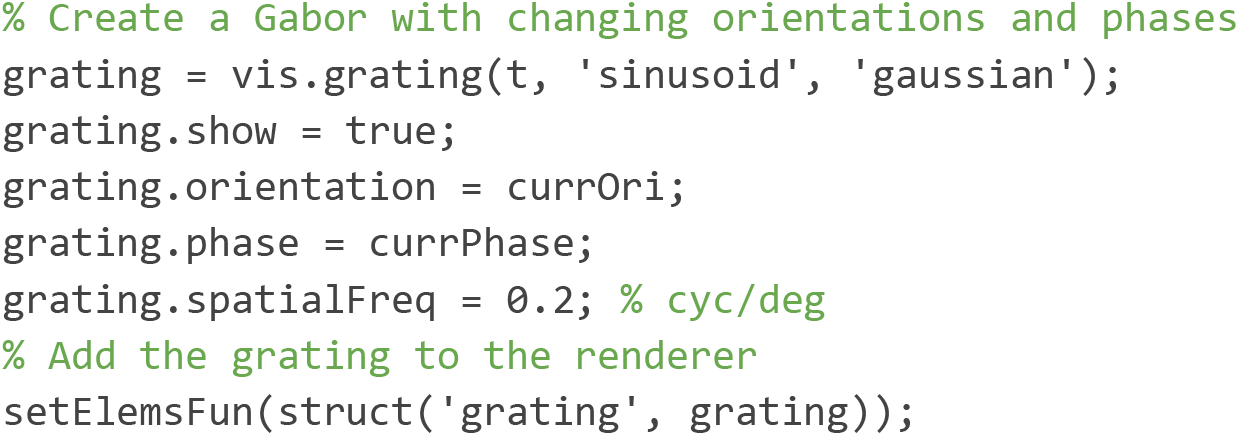

### Example 2: Pong

A second human-interactive Signals experiment contained in the Rigbox repository is an exp def which runs the classic computer game, Pong (Figure 8). The signal which sets the player’s paddle position is mapped to the optical mouse. The epoch structure is set so that a trial ends on a score, and the experiment ends when either the player or cpu reaches a target score. The code is divided into three sections: 1) initializing the game, 2) updating the game, 3) creating visual elements and defining exp def parameters. To run this exp def, follow the directions in the header of the signals/docs/examples/signalsPong.m file in the Rigbox repository. Because the file itself (including copious documentation) is over 300 lines, we will share only an overview here; however, readers are encouraged to look through the full file at their leisure.

**Figure 8:**
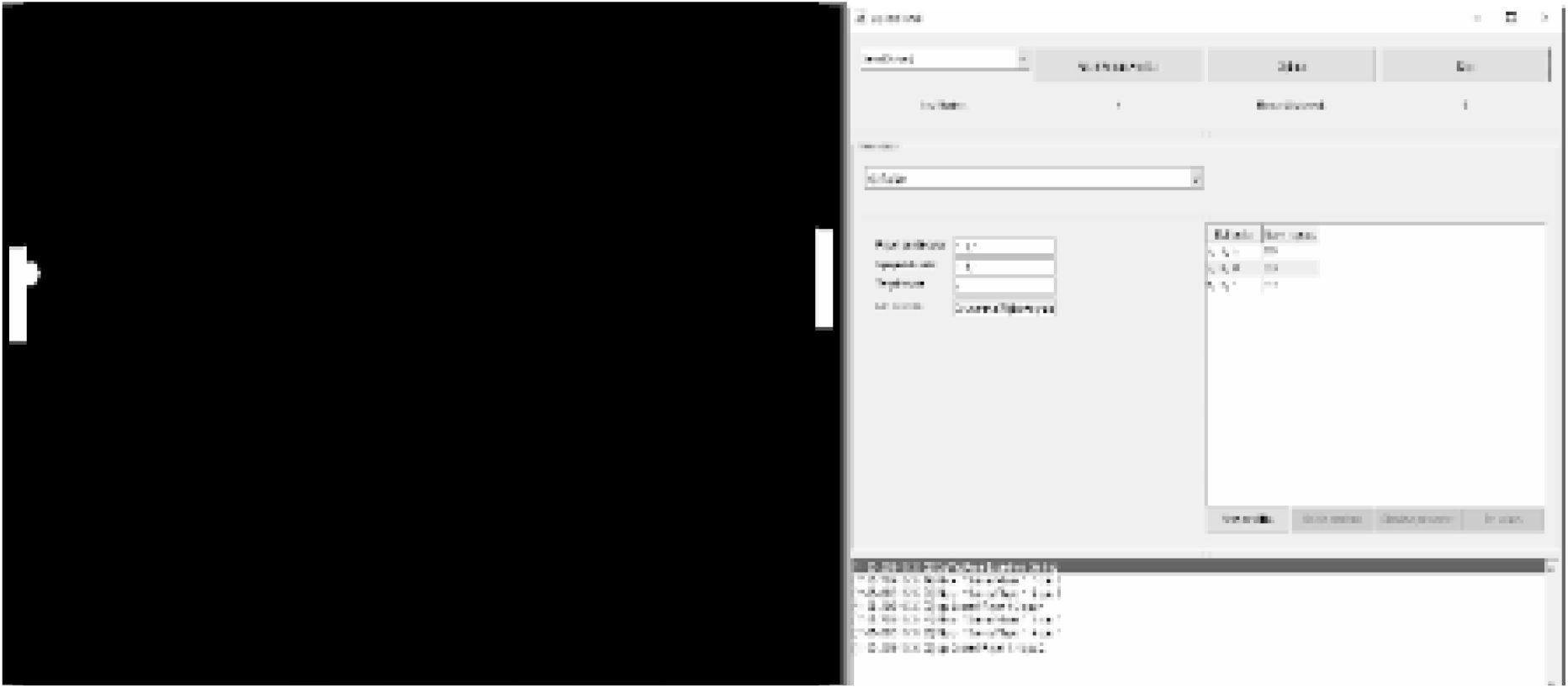
A screenshot of Pong run in Signals.

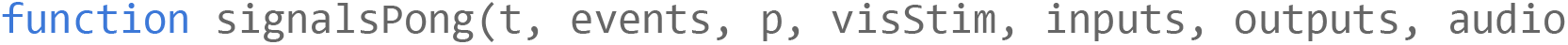

In this first section, we define constants for the game, arena, ball, and paddles:

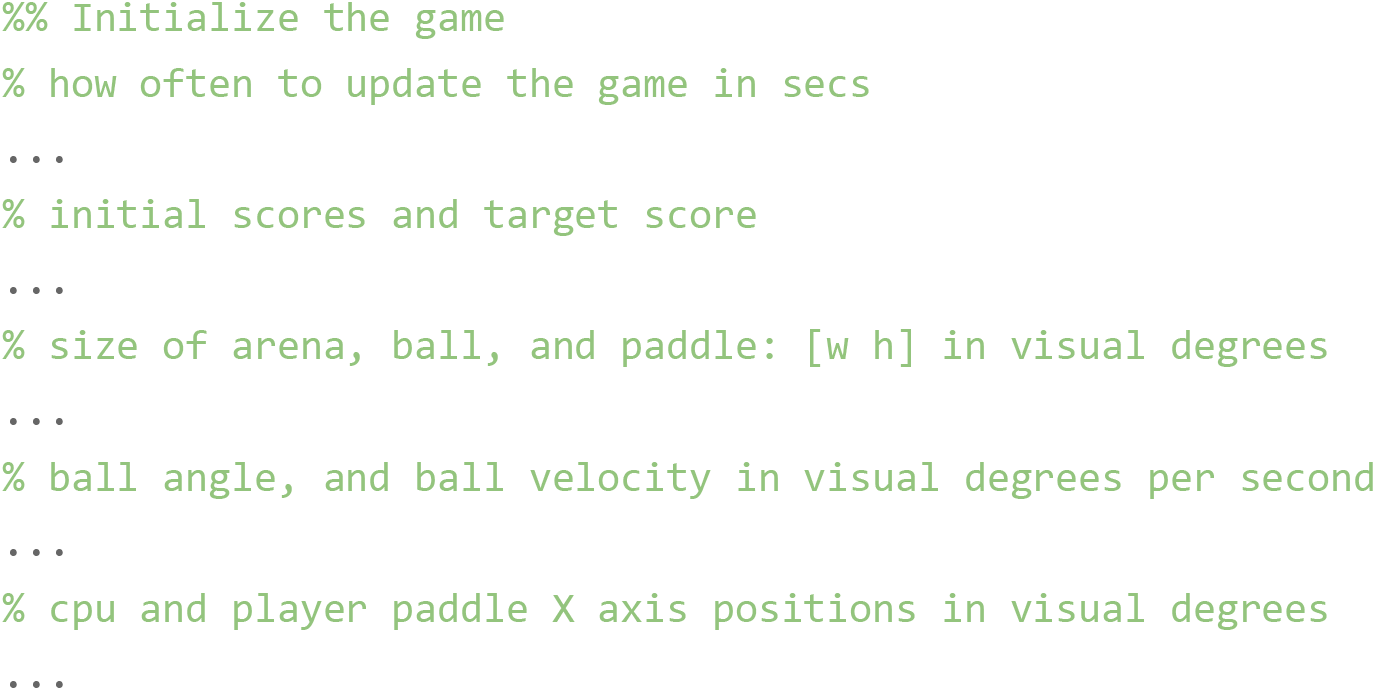

The helper function, getYPos, returns the y-position of the cursor, which will be used to set the player paddle:

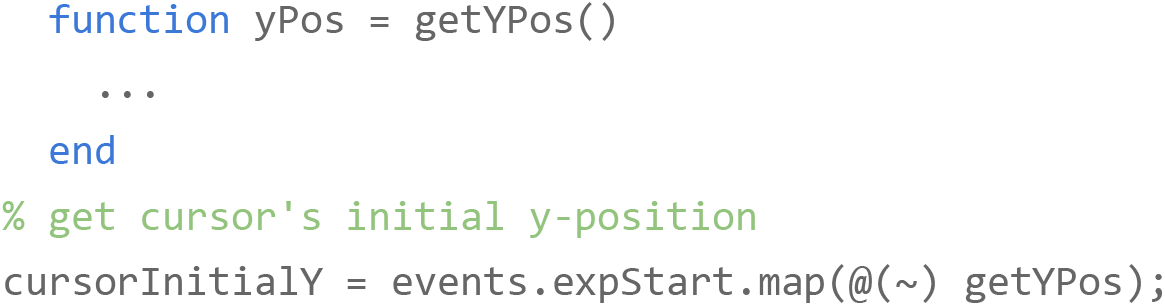

In the second section, we define how the ball and paddle interactions update the game:

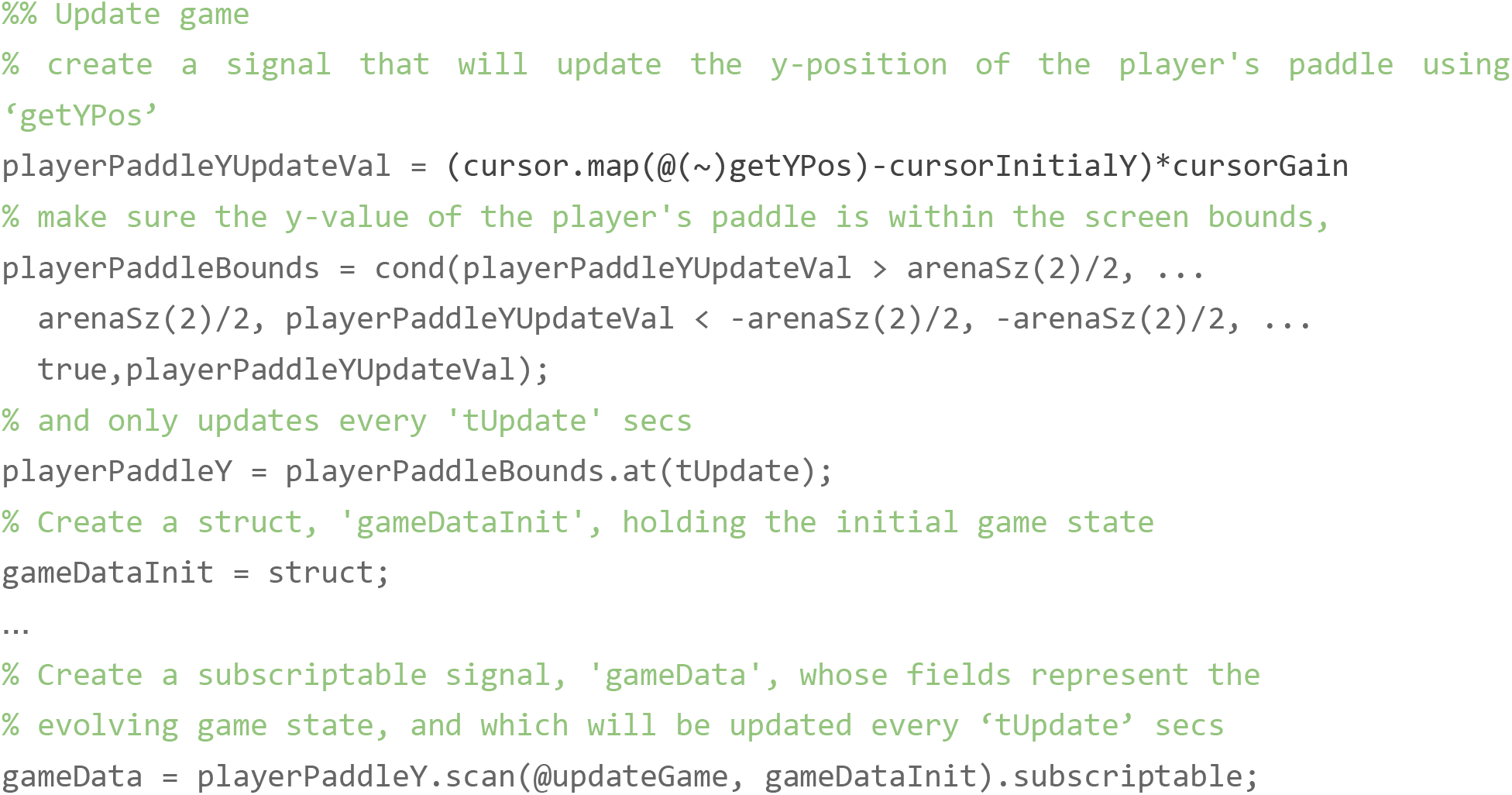

The helper function, updateGame, updates gameData. Specifically, it updates the ball angle, velocity, position, cpu paddle position, and player and cpu scores, based on the current ball position:

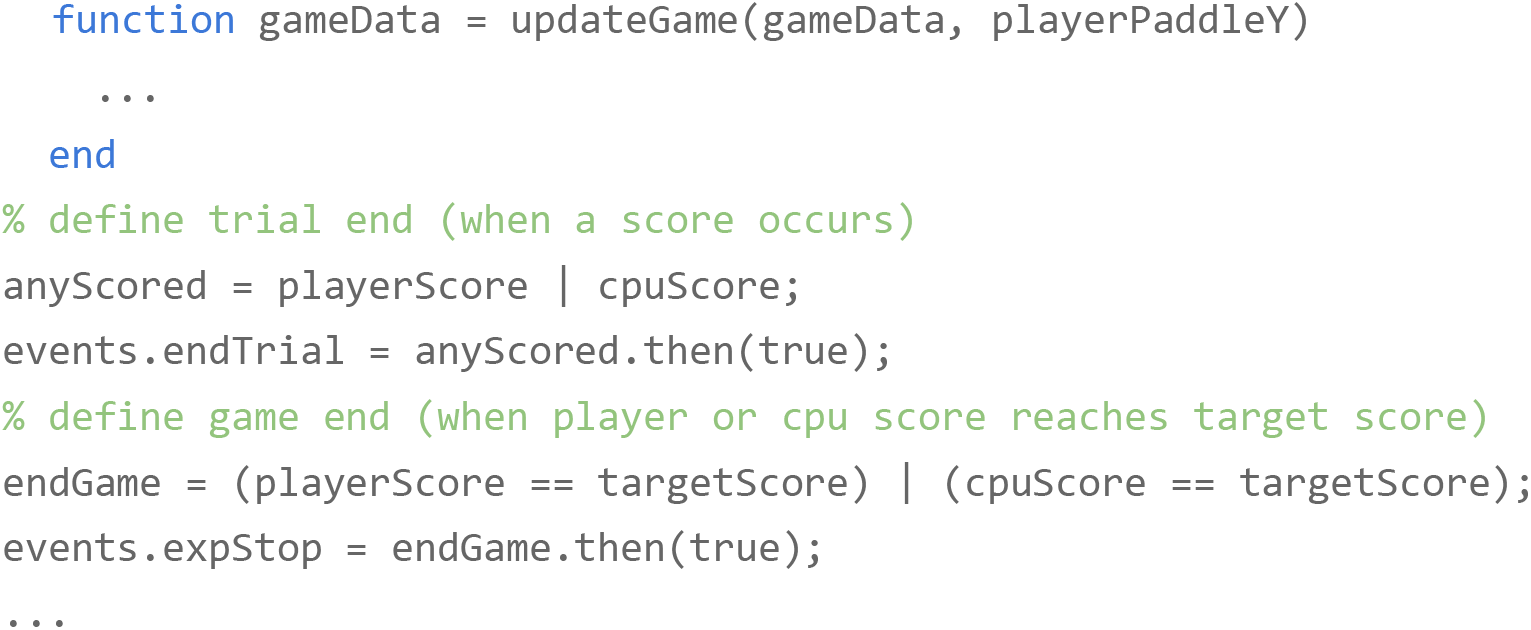

In the final section, we create the visual elements representing the arena, ball, and paddles, and define the exp def parameters:

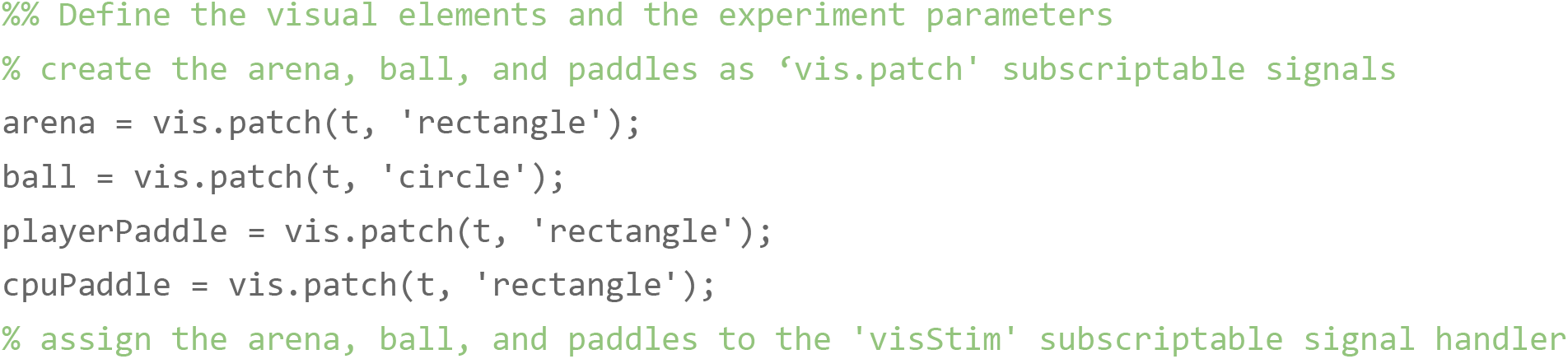
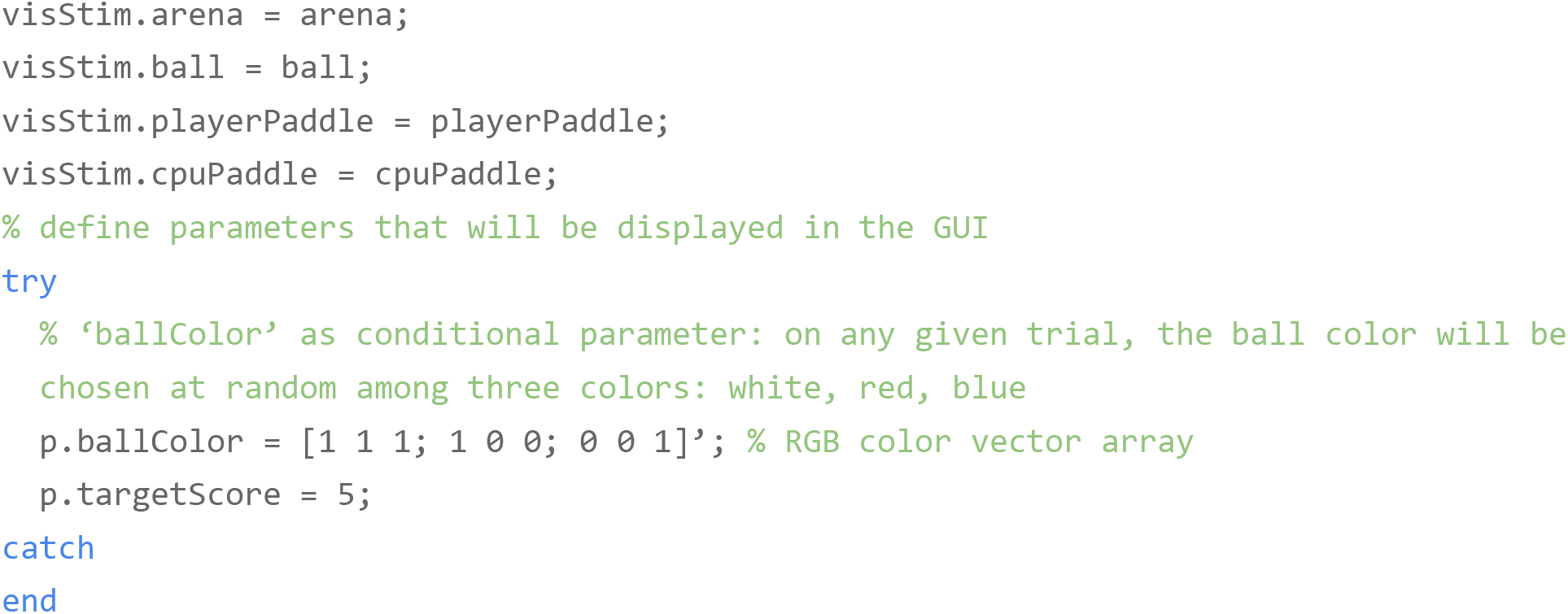

### Benchmarking

Fast execution of experiment runtime code is crucial for performing and accurately analyzing results from a behavioral experiment. Here we show benchmarking results for the Signals framework. We include results for individual operations on a signal and for operations which propagate through each signal in a network. Single built-in MATLAB operations and Signals-specific methods are consistently executed in the microsecond range (Figure 9). The network used in the Burgess Steering Wheel Task (`signals/docs/examples/advancedChoiceWorld.m`) contains 338 signals spread over 10 layers; a similar network of 350 signals spread over 20 layers can update all signals in under 5 milliseconds, and a network of 120 signals spread over 20 layers can update all signals with sub-millisecond precision (Figure 10). Lastly, we include results from running the Burgess Steering Wheel Task in Signals: updates of the wheel position typically took less than 2 milliseconds, the time between rendering and displaying the visual stimulus typically took less than 15 milliseconds, and the delay between triggering and delivering a reward was typically under 0.2 milliseconds (Figure 11).

**Figure 9:**
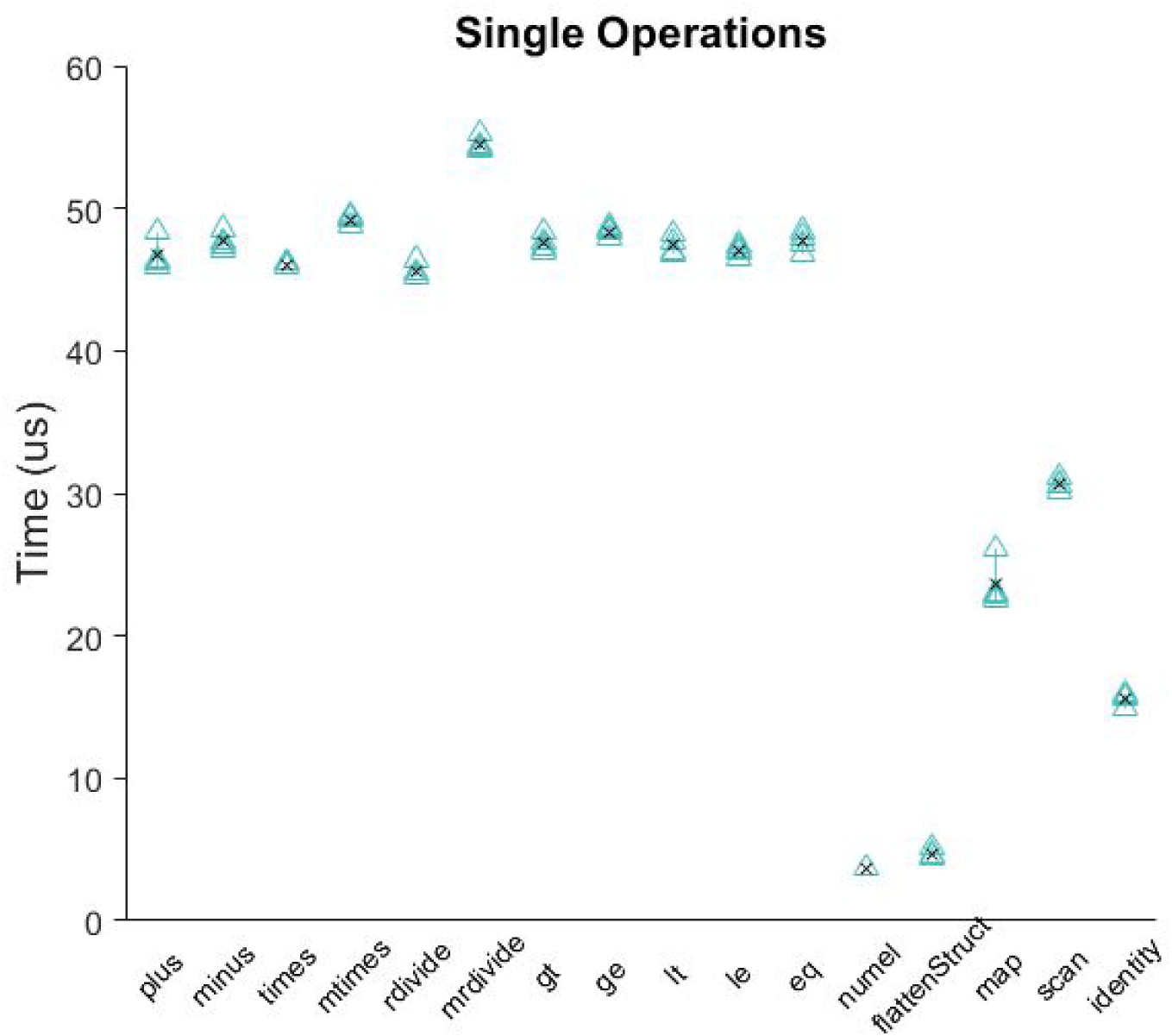
Benchmarking results for operations on a single signal. The black “x” shows the mean value per group.

**Figure 10:**
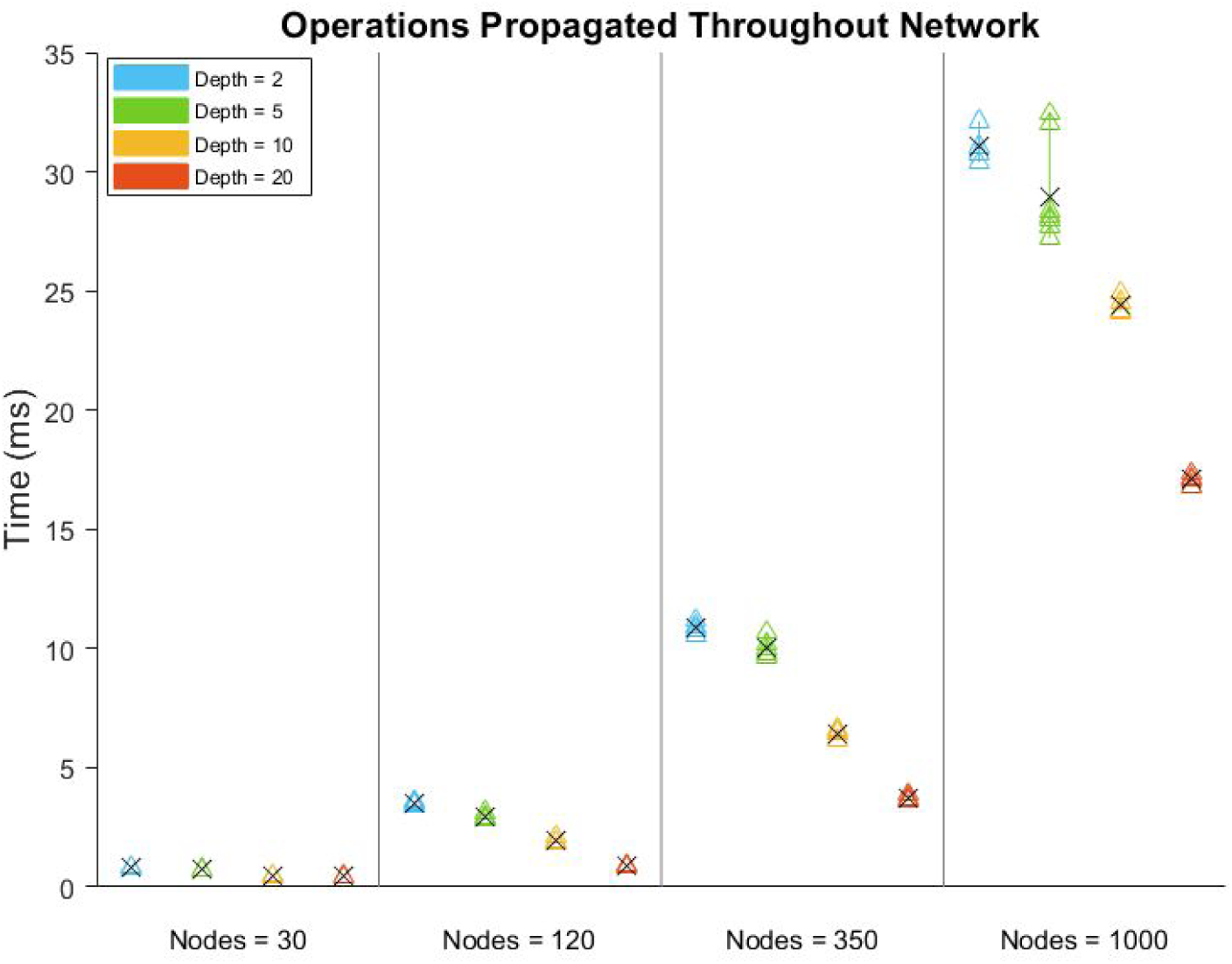
Benchmarking results for updating every signal in a network, for networks of various number of signals (nodes) spread over various number of layers (depth). The black “x” shows the mean value per group.

**Figure 11:**
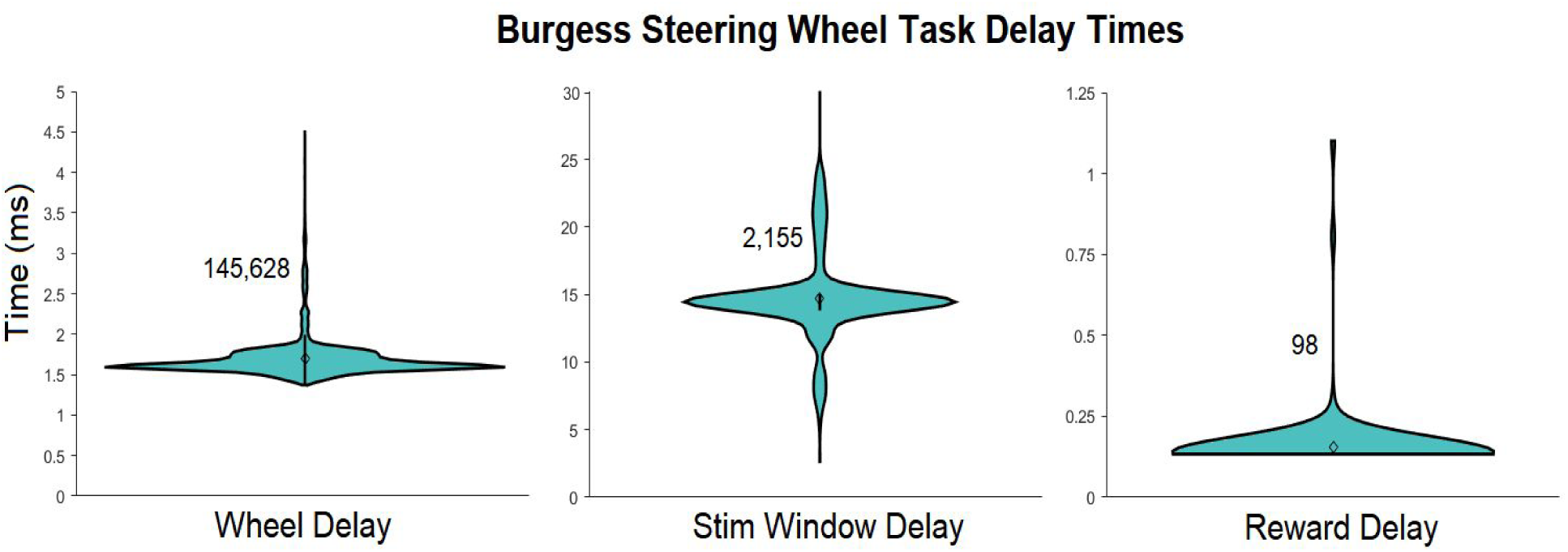
Delay times for specific updates when running the Steering Wheel Task. The number next to each violin plot indicates the number of samples in the group. “Wheel Delay” is the time between polling consecutive position values from the hardware wheel. “Stim Window Delay” is the time between triggering a display to be rendered, and it’s complete render on a screen. “Reward Delay” is the time between triggering a reward to be delivered and its delivery. 99th percentile outliers were not included in the plot for “Wheel Delay”: there were 98 instances in which the wheel delay took between 200-600 ms, due to execution time of the NI-DAQmx MATLAB package when sending analog output (reward delivery) via the USB-6211 DAQ.

All results in the Benchmarking section were obtained from running MATLAB 2018b on a Windows 10 64-bit OS with an Intel core i7 8700 processor and 16 GB DDR4 dual channel RAM clocking at a double data rate of 2133 MHz. Because single executions of signals operations were too quick for MATLAB to measure precisely, we repeated operations 1,000 times and divided MATLAB’s returned measured time by 1,000. MATLAB 2018b’s Performance Testing Framework was used to obtain these results. `signals/tests/Signals_perftest.m` contains the code used to generate the results shown in Figure 9. `signals/tests/results/2019-06-14_Signals_perftest.mat` contains a table of the data used to generate these results. `signals/tests/results/2019-06-04_advancedChoiceWorld_Block.mat` contains the data used to generate the results shown in Figure 10. National Instrument’s USB-6211 was used as the data acquisition device.

### The Other Packages in Rigbox

Often experiments are iterative: task parameters are added or modified many times over, and finding an ideal parameter set can be an arduous process. Rigbox allows the experimenter to develop and test their experiment without having to worry about boilerplate code and UI modifications, as these are handled by other packages in a modular fashion. Much of the code is object-oriented with most aspects of the system represented as configurable objects. Below is a short description of each package.

The **hardware package** *+hw* contains calibration functions and classes for interfacing with various hardware. These include abstract classes such as `Window` and `DataLogger`, which define general methods for high-level interaction with an on-screen stimulus display window and a hardware logging device, respectively. `Window` and `DataLogger` have concrete implementations for specific systems: `+ptb/Window` subclasses `Window` to represent a Psychophysics Toolbox stimulus window, and `DaqRotaryEncoder` subclasses `PositionSensor` (which subclasses `DataLogger`) to represent the Lego wheel used in the Burgess Steering Wheel Task (Figure 12). For novel implementations, additional subclasses can be created from these abstract classes to represent other specific hardware.

**Figure 12.**
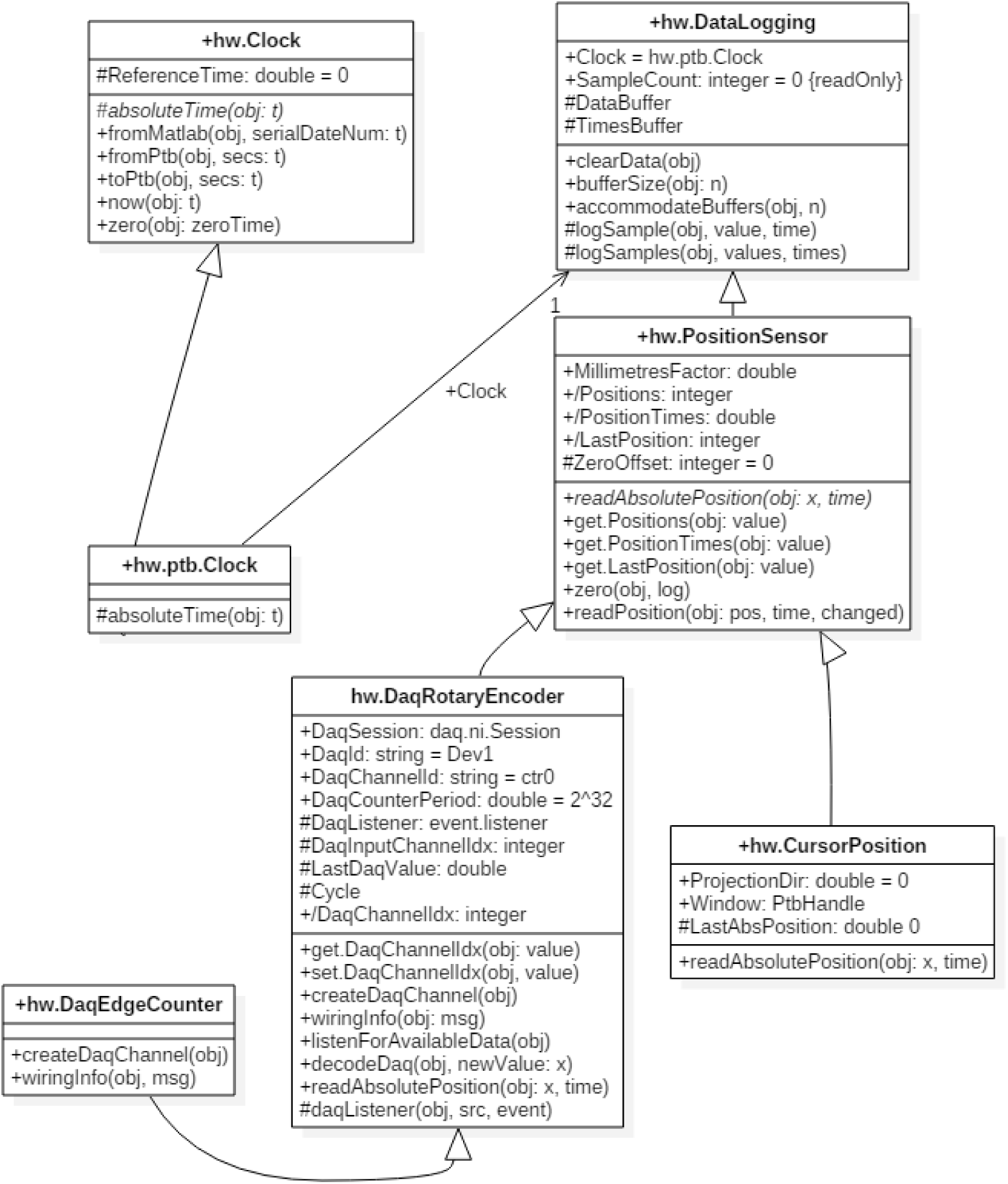
A UML diagram depicting the class structure for data logging in Rigbox. Each box represents a class and contained within it is the name, attributes and methods. The superclass is `DataLogging`, which contains the most general attributes and methods. `PositionSensor`, it’s immediate subclass (indicated by the white arrow) provides general abstract methods such as `readAbsolutePosition` for reading the raw position of some nondescript linear position sensor. The implementation of this depends on the specific drivers and hardware of each device. Two such subclasses are shown: one for interfacing with a rotary encoder via a NI-DAQ, and another for reading cursor position. The specific details of this need only be known to each subclass, and therefore it is straightforward to swap in different devices without having to modify other parts of the system. Also shown is the abstract Clock class and its concrete implementation using the Psychophysics Toolbox. The clock object is used in numerous different hardware classes and ensure that all run via the same clock.

The hardware package also contains a class called `Timeline` which manages the acquisition and generation of experimental timing data using a National Instruments Data Acquisition Device (NI-DAQ) (Figure 13). The main timing signal, chrono, is a digital square wave that flips each time a new chunk of data is available from the NI-DAQ. A callback function to this flip event collects the NI-DAQ timestamp of the scan where each flip occured. The difference between this timestamp and the system time recorded when the flip command was given is recorded as an offset time. This offset time can be used to unify all timestamps across computers during an experiment. Thus, all event timestamps across all computers for a given experiment are recorded in times relative to chrono. A `Timeline` object can acquire any number of hardware events and record their values with respect to this offset; for example, a `Timeline` object can record when a reward is delivered, a laser is fired, a sensor is interacted with, a screen displaying visual stimuli is updated, etc. In addition to chrono, a `Timeline` object can also output TTL and clock pulses for triggering external devices (e.g. to acquire frames at a specific rate).

**Figure 13.**
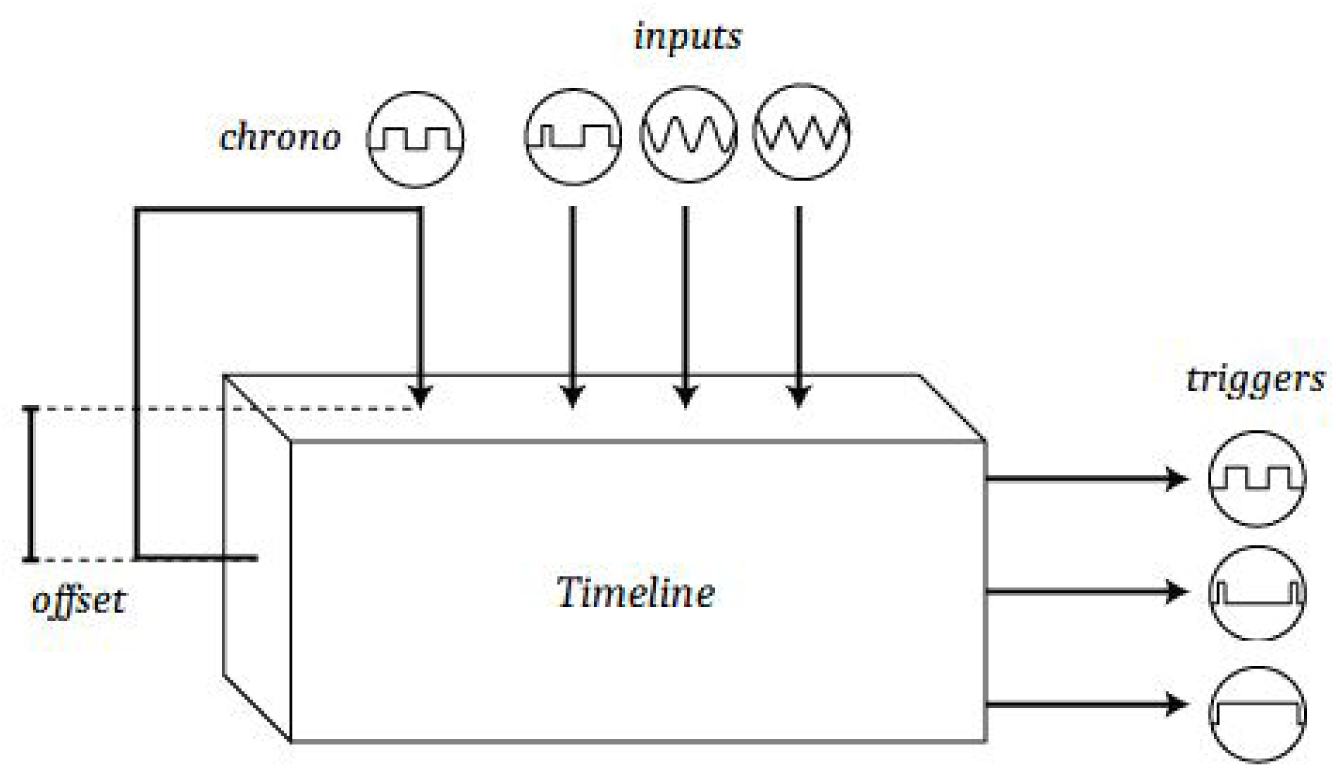
Representation of a Timeline object. The top most signal is the main timing signal, “chrono”, which is used to unify all timestamps across computers during an experiment. The “inputs” represent different hardware input signals read by a NI-DAQ, and the “triggers” represent different hardware output signals, triggered by a NI-DAQ.

The **data package** *+dat* contains a number of simple functions for saving and locating data. Data organization supports separation of data types between repositories, and redundant local and remote storage. Because all code uses the same paths file, it is very simple to change the location of data and configuration files. Furthermore, this system can be easily used with one’s own code to generate read and write paths for arbitrary datasets. The **experiment package** *+exp* contains all of the code pertaining to experiment setup and configuration. Two key aspects of this are the `Parameters` class, which sets, validates, and assorts experiment conditions for each experiment, and the `SignalsExp` class, which runs an experiment after loading in the experimenter’s exp def and appropriate parameters into a Signals network.

The **server package** *+srv* provides high level network communication between MC and SC. In addition, this package provides functions for triggering remote recording software via UDPs.

The **experiment UI package** *+eui* provides all the graphical user interface (GUI) code. Principally, this is employed for the `mc` function, which launches the main GUI on MC. The MC GUI is used to load and configure experiment parameters on MC, monitor experiments through customizable plots, view experiment history, and log metadata.

The **psychometrics package** *+psy* contains simple functions for processing and plotting psychometric data.

The **alyx-matlab** package serves as a MATLAB client for interfacing with an Alyx database. This package allows experimenters to make queries and posts to an Alyx database within MATLAB, and create notes during an experiment which are automatically synced to the database. alyx-matlab uses the npy-matlab submodule to provide support for saving data.

Alyx is a meta-database that allows experimenters to keep track of animal procedures, such as breeding and implantation, and organize experimental sessions and their associated files (Rossant et. al, 2018). The database is heavily used by the International Brain Laboratory due to its lightweight nature, and can be easily installed on most web servers (Abbott et al., 2017). More information on Alyx and alyx-matlab can be found in `alyx-matlab/docs` within the Rigbox repository. The use of Alyx and alyx-matlab within Rigbox is optional.

## Discussion

In our laboratory, Rigbox is at the core of our operant, passive, and conditioning experiments. The principal behavioral task we use is the Burgess Steering Wheel Task (Burgess et al., 2017). Using Rigbox, we have been able to create multiple variants of this task. These have included unforced choice, multisensory choice, behavior matching, and bandit tasks, using wheels, levers, balls, and lick detectors. In addition, Rigbox has allowed us to rapidly integrate these tasks with a variety of recording techniques, including electrode recordings, 2-Photon imaging, and fiber photometry, and neural perturbations, such as scanning laser inactivation and dopaminergic stimulation (Jun et al., 2017; Jacobs et al., 2018; Lak et al., 2018; Steinmetz et al., 2018; Shimaoka et al., 2018; Zatka-Haas et al., 2018).

Given the modular nature of Rigbox, new features and hardware support may be easily added, provided there is driver support in MATLAB. For example, to add support for a novel data acquisition device (such as an Arduino or other microcontroller), one can simply create a subclass of the `+hw/DaqController` class. Similarly, to add support for a novel position sensor, a new `+hw/PositionSensor` subclass could be created. These classes simply define what happens when, for example, the code triggers a hardware output, or polls a hardware input. This principle also holds true for implementing various visual stimulus viewing models, of which there is currently only one. A new viewing model class could be implemented to allow for virtual reality experiments, for example.

To the best of our knowledge, Rigbox is the most complete behavioral control software toolbox currently available in the neuroscience community; however, a number of other toolboxes implement similar features in different ways (Bcontrol 2014; Sanders 2019; Akam 2019; Aronov and Tank, 2014) (Table 1). Some of these toolboxes also include some features not currently available in Rigbox, for example microsecond precision triggering of within-trial events and creating 3D virtual environments. Indeed, the features employed by a particular toolbox have advantages (and disadvantages) depending on the user’s desired experiment.

**Table 1:** Comparison of major features across behavioral control system toolboxes. The top row contains the toolbox names, and the first column contains information on a feature’s implementation. Note: the toolboxes and features mentioned in this table are not exhaustive.

|  | **BControl** | **Bpod** | **pyControl** | **VirMEN** | **Rigbox** |
| --- | --- | --- | --- | --- | --- |
| Behavioral task design paradigm | Procedural | Procedural | Procedural | Object-Oriented | Functional Reactive |
| Implements viewing model? 3D viewing model? | no | no | no | yes, yes | yes, no |
| Interfaces with hardware? | yes | yes | yes | yes | yes |
| Synchronizes multiple datastreams? | yes | yes | yes | no | yes |
| Communicates with a remote database? | yes | yes | no | no | yes |

There are pros and cons to following different programming paradigms for software developers who decide how a user will programmatically design a behavioral task. Generally, three main paradigms exist: procedural, object-oriented, and functional reactive. Here, in the context of programmatic task design, we briefly discuss the differences between these paradigms and in which scenarios one may be favored over the others. Note: here we only discuss the aspect of a toolbox that deals with behavioral task design, not the overall structure of a toolbox (e.g. Rigbox is built on an object-oriented paradigm, but Signals provides a functional reactive paradigm in which to implement a behavioral task).

A procedural approach to task design is probably the most familiar to behavioral neuroscientists. This approach focuses on “how to execute” a task by explicitly defining a control flow that moves a task from one state to the next. The Bcontrol, Bpod, and pyControl toolboxes follow this paradigm by using a real-time finite state machine (RTFSM) which controls a task’s state (e.g. initial state, reward, punishment, etc.) during each trial. Some advantages of this approach are that it’s simple and intuitive, and guarantees event timing precision down to the minimum cycle of the state machine (e.g. Bcontrol RTFSMs run at a minimum cycle of 6 KHz). Some disadvantages of this approach are that the memory for task parameters are limited by the RTFSM’s number of states, and that the discrete implementation of states isn’t amenable to experiments which seek to control parameters continuously (e.g. a task which uses continuous hardware input signals).

Like the procedural approach to task design, an object-oriented approach also tends to be intuitive: objects can neatly represent an experiment’s state via datafields. Objects representing experimental parameters can easily pass information to each other, and trigger experimental states via event callbacks. The VirMEn toolbox implements this approach by treating everything in the virtual environment as an object, and having a runtime function update the environment by performing method calls on the objects based on input sensor signals from a subject performing a task. Some disadvantages of this approach are that the speed of experimental parameter updates are limited by the speed at which the programming language performs dynamic binding (which is often much slower than the RTFSM approach discussed above), and that operation “side effects” (which can alter an experiment’s state in unintended ways) are more likely to occur due to the emphasis on mutability, when compared to a pure procedural or functional reactive approach.

By contrast, Signals follows a functional reactive approach to task design. As we have seen, some advantages of this approach include simplifying the process of updating experimental parameters over time, endowing parameters with memory, and facilitating discrete and continuous event updates with equal ease. In general, a task specification in this paradigm is declarative, which can often make it clearer and more concise than in other paradigms, where control flow and event handling code can obscure the semantics of the task. Some disadvantages are that it suffers from similar speed limitations as in an object-oriented approach, and programmatically designing a task in a functional reactive paradigm is probably unfamiliar to most behavioral neuroscientists. When considering the entire set of behavioral tasks, no single programming paradigm is perfect, and it is therefore important for an experimenter to consider the goals for their task’s implementation accordingly.

Rigbox is currently under active, test-driven development. All our code is open source, distributed under the Apache 2.0 license, and we encourage users to contribute. Please see the contributing guidelines in the repository for contributing code and reporting issues.

## Acknowledgments

We thank Nick Steinmetz, Max Hunter, Peter Zatka-Haas, Kevin Miller, Hamish Forrest, and other members of the lab for troubleshooting, feedback, inspiration, and code contribution. This work was funded by the Medical Research Council (Doctoral Training Award to CPB), the Royal Society (Newton International Fellowship to AJP), EMBO (fellowship to AJP), the Human Frontier Science Program (fellowship to AJP), and by the Wellcome Trust (grant 205093 to MC and KDH). MC holds the GlaxoSmithKline / Fight for Sight Chair in Visual Neuroscience.

